# Peptide modified, programmable DNA tetrahedra to modulate autophagy in biological systems

**DOI:** 10.1101/2024.11.03.621781

**Authors:** A Hema Naveena, Krupa Kansara, Nihal Singh, Sharad Gupta, Ashutosh Kumar, Dhiraj Bhatia

## Abstract

Autophagy is a critical cellular pathway for degrading and recycling damaged components, essential for maintaining cellular homeostasis. Dysregulation of autophagy contributes to various diseases, including neurodegenerative disorders, cancers, and metabolic syndromes, highlighting the therapeutic potential of controlled autophagy induction. However, current autophagy inducers often lack specificity and may inadvertently trigger apoptosis, limiting their clinical utility. Here, we present a DNA tetrahedron-BH3 peptide nanosystem (Tdpep) engineered to selectively induce autophagy by disrupting the Beclin 1-Bcl2 interaction, a pivotal regulatory point in autophagy initiation. Tdpep, functionalized with a BH3 peptide targeting Bcl2, demonstrated efficient cellular uptake and minimal cytotoxicity in HeLa cells at concentrations up to 200nM. Autophagy induction was confirmed by increased LC3B puncta formation and fluorescence intensity comparable to that induced by rapamycin. Autophagy flux analysis of Tdpep with bafilomycin A1 validated enhanced autophagic activity rather than flux inhibition. Furthermore, Tdpep treatment significantly reduced cellular ROS levels, indicating effective autophagic turnover. Apoptosis assays showed that Tdpep did not induce apoptosis, confirming its selective autophagy induction. Furthermore, Tdpep nanosystem also induced autophagy in *Danio rerio* larvae in vivo model. Thus, this targeted DNA tetrahedron nanosystem provides a precise autophagy modulation platform with minimized off-target effects, offering a promising therapeutic strategy for diseases associated with autophagy dysfunction.

**Graphical abstract:** 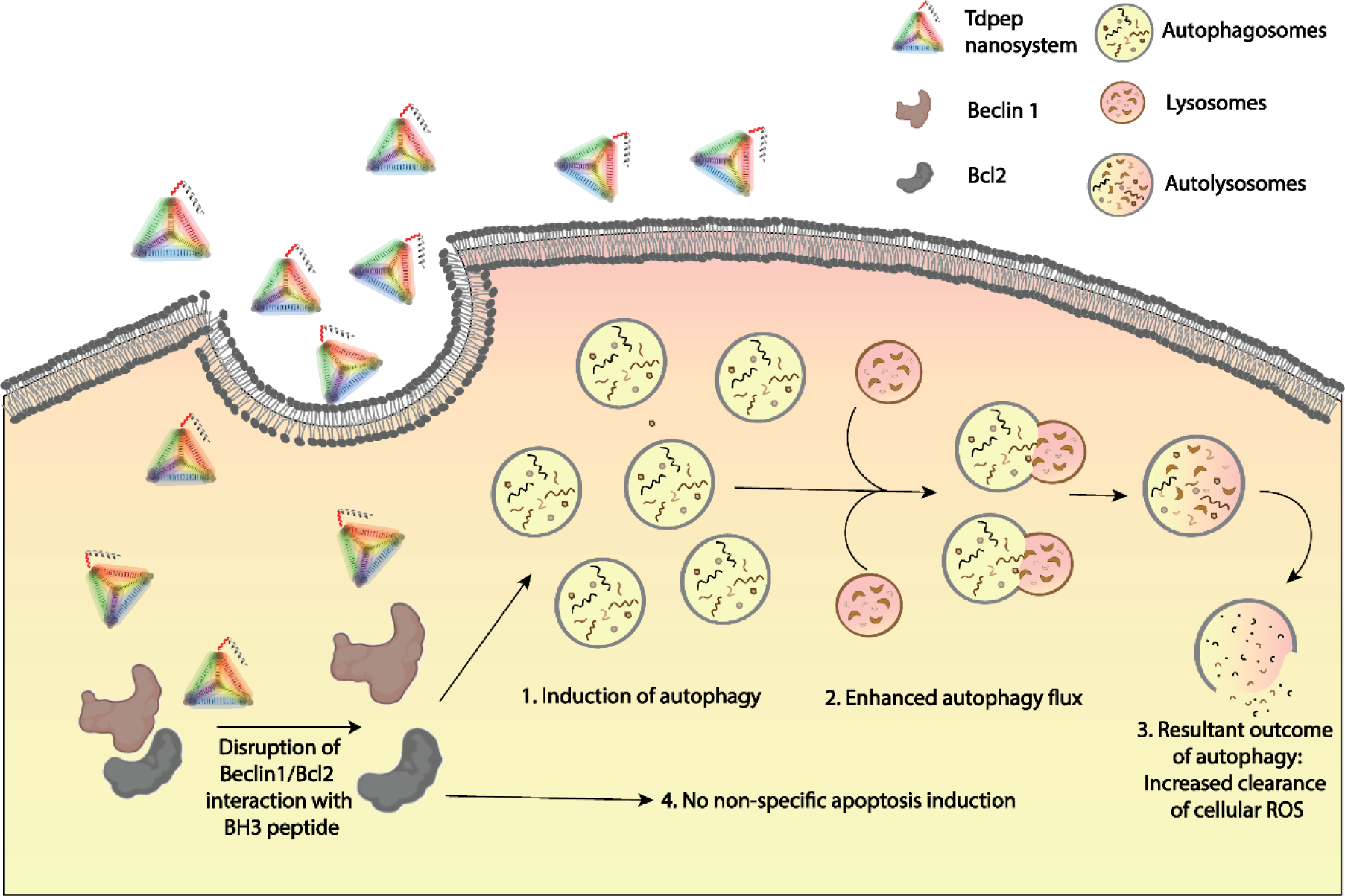

## 1. Introduction

Macroautophagy (referred as autophagy hereafter) is a fundamental and conserved cellular process through which cells degrade and recycle their own constituents (1). This self-degradative mechanism involves the formation of double-membrane vesicles, termed autophagosomes, that engulf cellular components, which are then fused with lysosomes for degradation and recycling. Autophagy plays a crucial role in maintaining cellular homeostasis by clearing damaged organelles(2), aggregated proteins(3), and intracellular pathogens (4–6) . By maintaining a balance between synthesis, degradation, and recycling, autophagy supports cellular health and function under both normal and stress conditions(1).

Modulation of autophagy is increasingly recognized for its potential to impact disease pathology(7). Dysregulation of autophagy has been implicated in a wide variety of diseases, including neurodegenerative disorders (e.g., Alzheimer’s(8) and Parkinson’s diseases (9)), cardiovascular diseases (10), metabolic disorders (11,12), and various cancers (13–15). In neurodegenerative diseases, for example, impaired autophagy contributes to the accumulation of toxic protein aggregates (8,9), whereas in cancer, autophagy may act as a double-edged sword, either suppressing or promoting tumor progression depending on the cellular context (13–15). Thus, controlled induction of autophagy presents a promising therapeutic avenue for managing diseases characterized by autophagy dysregulation (16). By enhancing autophagic activity, cells may clear disease-causing proteins, damaged organelles, or pathogens more effectively, potentially preventing disease onset, progression, or exacerbation (7,16).

To date, a variety of strategies have been employed to induce autophagy, ranging from pharmacological agents to genetic modulation(16). Although effective, these strategies often lack specificity, impacting autophagy indiscriminately across different cell types and tissues. Moreover, broad-spectrum autophagy induction can lead to off-target effects, complicating the clinical application of such treatments due to the potential for unintended cellular stress or damage (7). Moreover, most nanoparticle-based approaches inadvertently induce apoptosis alongside autophagy, with limited clarity on the exact mechanisms by which they influence autophagy pathways (17–19).

In an effort to overcome these limitations, we have developed a DNA tetrahedron- peptide based nano system designed for targeted autophagy induction via selective disruption of the Beclin 1- Bcl2 complex. The Beclin 1-Bcl2 complex is a key regulatory checkpoint in autophagy initiation, where Bcl2, an anti-apoptotic protein, inhibits autophagy by binding to Beclin 1, an essential autophagy activator. Disruption of this complex frees Beclin 1 to initiate autophagy, making it a precise target for controlled autophagy induction (20). In this study, we designed a BH3 peptide that selectively disrupts the interaction between Beclin 1 and Bcl2 to induce autophagy. To enhance cellular uptake, and to further enable tissue specific targeting, the peptide was conjugated to a DNA tetrahedron nanostructure. The DNA tetrahedron enhances the peptide’s cellular internalization, and allows for modular customization to achieve spatially precise targeting (21–23).

The devdeloped peptide-functionalized DNA tetrahedron nanosystem provides a more selective and controlled strategy for autophagy induction, offering potential therapeutic applications in diseases associated with autophagy dysregulation, such as neurodegenerative disorders and cancers. Our approach is designed to reduce off-target effects common in broader-spectrum autophagy inducers, thus advancing the development of safer and more effective autophagy-modulating therapies.

## 2. Results and discussion

### 2.1 Synthesis of DNA tetrahedron-peptide nanosystem for induction of bulk autophagy

To induce autophagy, we designed a 21-amino acid BH3 peptide based on residues 108-123 (TMENLSRRLKVTGDLFDIMS) of Beclin1, a key region involved in the Beclin1-Bcl2 interaction. The Beclin1-Bcl2 complex is a regulatory hub for autophagy suppression, and disrupting this interaction has been shown to promote autophagy. To facilitate peptide coupling, an additional cysteine residue was introduced at the N-terminus of the peptide sequence. The engineered BH3 peptide can competes with Beclin1 for binding to Bcl2, thereby freeing Beclin1 to initiate autophagy (**Figure 1a**).

**Figure 1:**
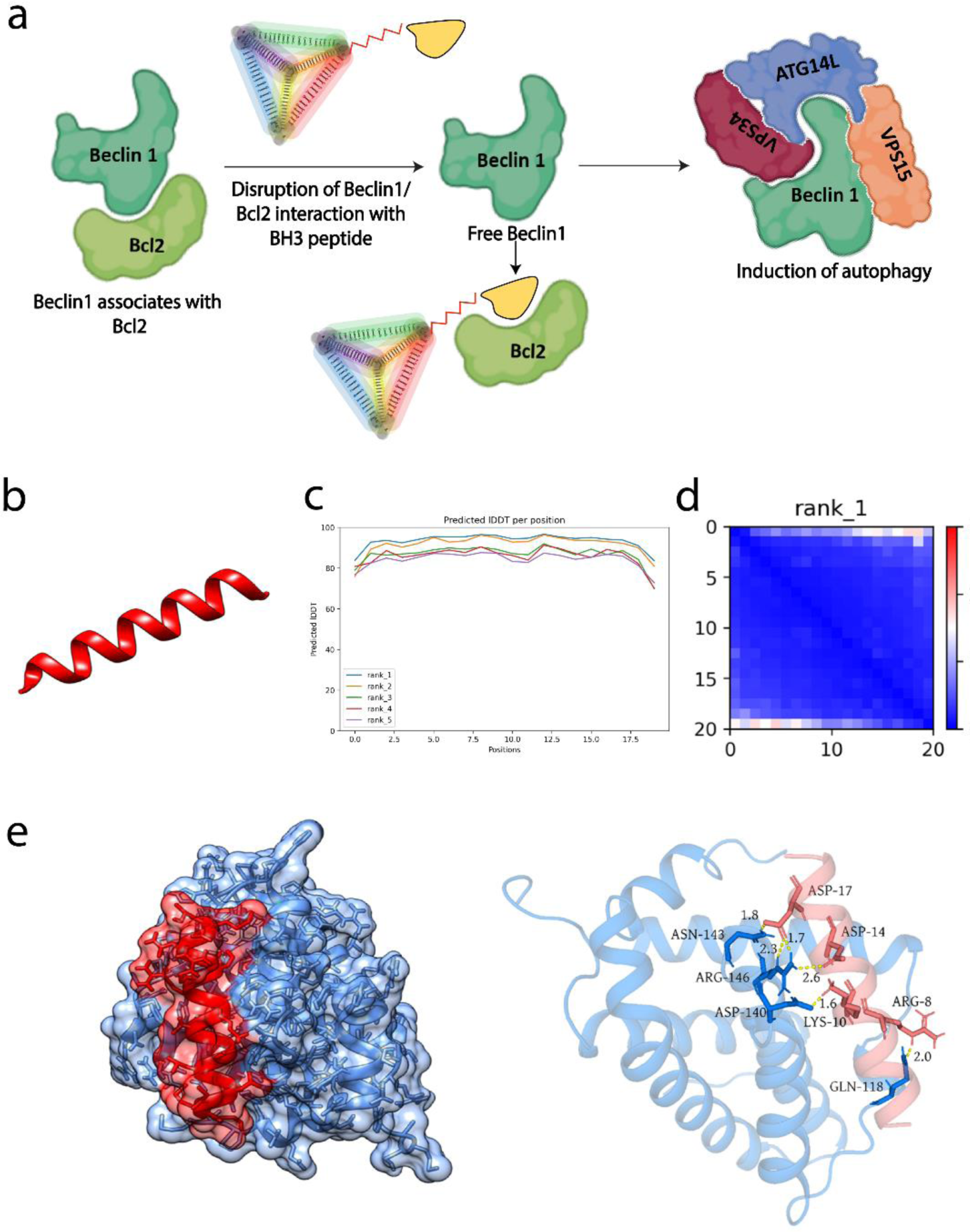
a. The schematics of the beclin1/bcl2 disruption and induction of autophagy with DNA tetrahedron BH3 peptide nanosystem. b. The predicted structure of BH3 peptide with AlphaFold 2.2. c. Predicted local distance difference test for the predicted BH3 peptide model. d. Predicted aligned error for the predicted BH3 peptide model. e. Docking of BH3 peptide and Bcl2 protein via Haddock 2.4

To characterize the structural properties of the designed BH3 peptide, we utilized AlphaFold 2.2 to predict its tertiary structure. The prediction yielded a high-confidence model with a predicted local distance difference test (pLDDT) score exceeding 90%, indicating structural reliability (**Figure 1b, c**). Additionally, the predicted aligned error (PAE) scores were consistently low, further supporting the accuracy of the peptide model across different structural regions (**Figure 1d**). Molecular docking simulations were then performed using Haddock 2.4, employing the AlphaFold-predicted BH3 peptide model and an experimentally resolved structure of Bcl2 (5VAU). The docking analysis revealed several interaction clusters, which were evaluated based on cluster size, root mean square deviation (RMSD), and Haddock scores (**Supporting information, Table 2**). Results indicated that the BH3 peptide binds specifically within the Beclin1-binding pocket of Bcl2 (**Figure 1e**), supporting its potential to disrupt the Beclin1-Bcl2 interaction and thereby induce autophagy.

The BH3 peptide was synthesized in-house via solid-phase peptide synthesis. The expected molecular mass for the full-length peptide sequence, CTMENLSRRLKVTGDLFDIMS, was calculated to be 2427.22 Da, and matrix-assisted laser desorption/ionization (MALDI) mass spectrometry analysis confirmed the mass of the synthesized peptide as 2427.72 Da, verifying identity of the synthesized peptide (**Figure 2a**).

**Figure 2.**
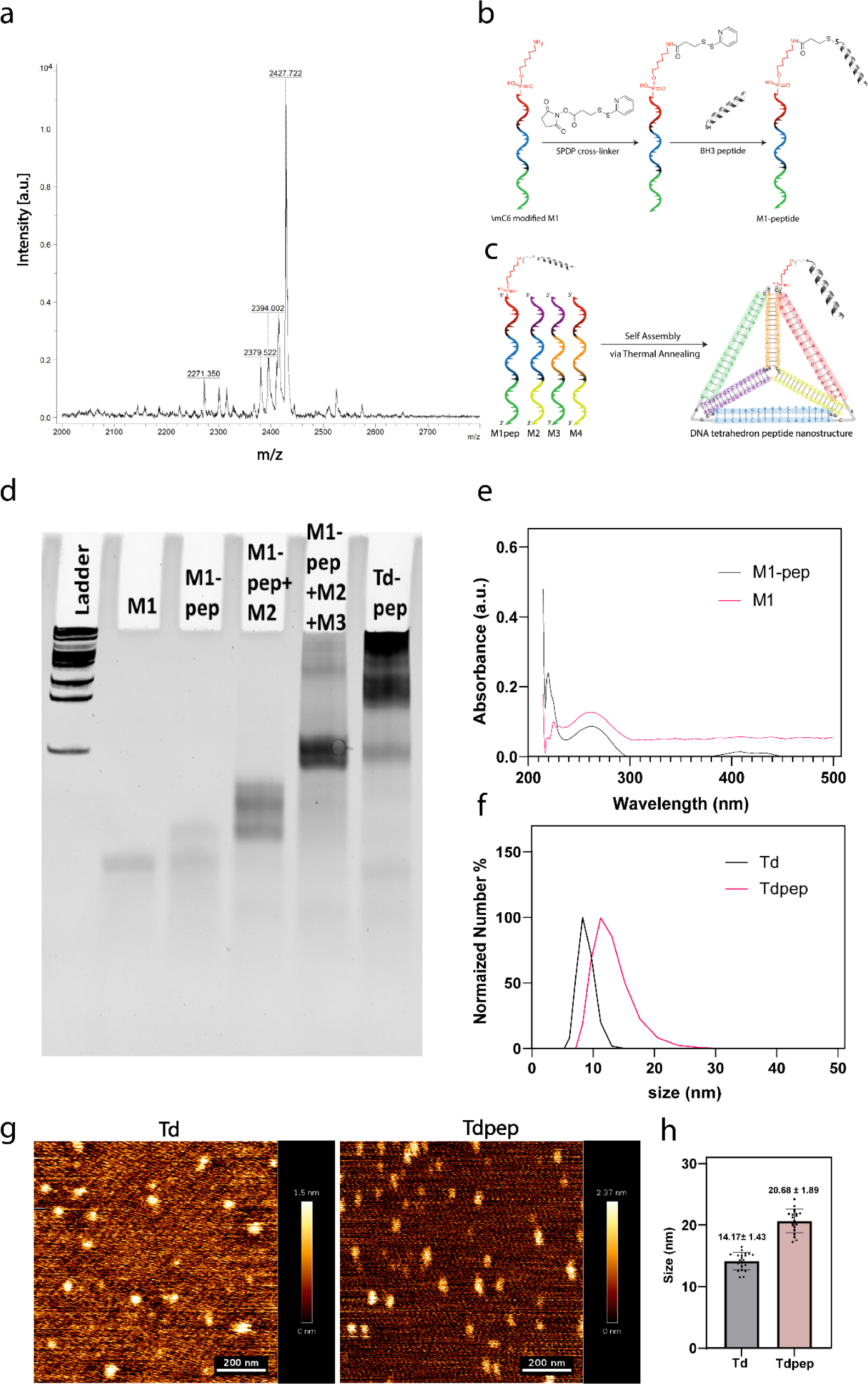
Synthesis and characterization of DNA Tdpep nanosystem. a. MALDI spectra of the synthesized BH3 peptide. b. Schematics of the SPDP coupling of the M1 oligonucleotide and the BH3 peptide. c. Schematics of the synthesis BH3 peptide functionalized DNA tetrahedron nanosystem (Tdpep). d. EMSA showing the gel based retardation in the 10% Native PAGE. e. UV-Visible spectrum of the M1 and M1-pep, showing the coupling of peptide with M1. f. DLS spectra of Td and Tdpep nanosystems. g. AFM images of the Td and Tdpep. Scale bar represents 200nm. h. Quantification of the size based on the AFM imaging of the Td and Tdpep nanosystems. Histobars represents mean ± SD of 20 particles.

For efficient cellular uptake, the BH3 peptide was conjugated to a DNA tetrahedron nanostructure (Td), which has been demonstrated to enhance cellular uptake due to its structural flexibility and amenability to functionalization. To achieve this, the peptide was coupled with the M1 oligonucleotide strand of the DNA tetrahedron using the SPDP (N-succinimidyl 3-(2- pyridyldithio) propionate) coupling chemistry, forming the M1-peptide (M1-pep) conjugate (**Figure 2b**). Successful coupling was confirmed through electrophoretic mobility shift assay (EMSA), as evidenced by a distinct band shift of the M1 strand (17059 Da) to M1-pep (19486 Da) (**Figure 2d**). UV-visible spectroscopy further validated the coupling, with the M1 oligonucleotide exhibiting an absorbance peak at ∼260 nm, and the M1-pep conjugate showing dual absorbance peaks at ∼260 nm and ∼220 nm, corresponding to the DNA and peptide components, respectively (**Figure 2e**).

The BH3 peptide-functionalized DNA tetrahedron nanodevice (Tdpep) was then synthesized in a one-pot thermal annealing reaction with four oligonucleotide strands (**Figure 2c**) (M1-pep, M2, M3, and M4; **Supplementary information, Table 3**). Formation of the Tdpep nanosystem was confirmed through several analytical techniques. EMSA displayed the characteristic ladder-like banding pattern of the DNA tetrahedron, indicative of higher-order structure formation (**Figure 2d**). Dynamic light scattering (DLS) measurements estimated the average hydrodynamic size of Td and Tdpep to be approximately 9 nm and 11 nm, respectively (**Figure 2f**). Atomic force microscopy (AFM) imaging further confirmed the size of Td and Tdpep, revealing nanosystems with an average diameter of ∼14 nm for Td and ∼20 nm for Tdpep, validating successful nanodevice synthesis and peptide conjugation (**Figure 2g, 2h**). These results collectively confirm the successful design and characterization of the BH3 peptide-conjugated DNA tetrahedron.

### 2.2 Tdpep nanosystem induces autophagy invitro

To evaluate the cellular uptake efficiency of the Td and Tdpep nanosystems in HeLa cells, the M4 oligonucleotide of each nanosystem was labeled with the Cy3 fluorophore. Uptake of the nanosystems was assessed at 30 minutes, 1 hour, and 2 hours’ post-treatment. Results indicated significant internalization of both Td and Tdpep nanosystems at all-time points, with enhanced uptake of the Tdpep system compared to Td alone across all intervals (**Figure 3a, 3b, 3c, 3d**). This enhanced internalization of the Tdpep suggests that the peptide conjugation may improve the uptake dynamics, however, the increased cellular internalization of the Tdpep system remains mechanistically unexplained at this stage.

**Figure 3:**
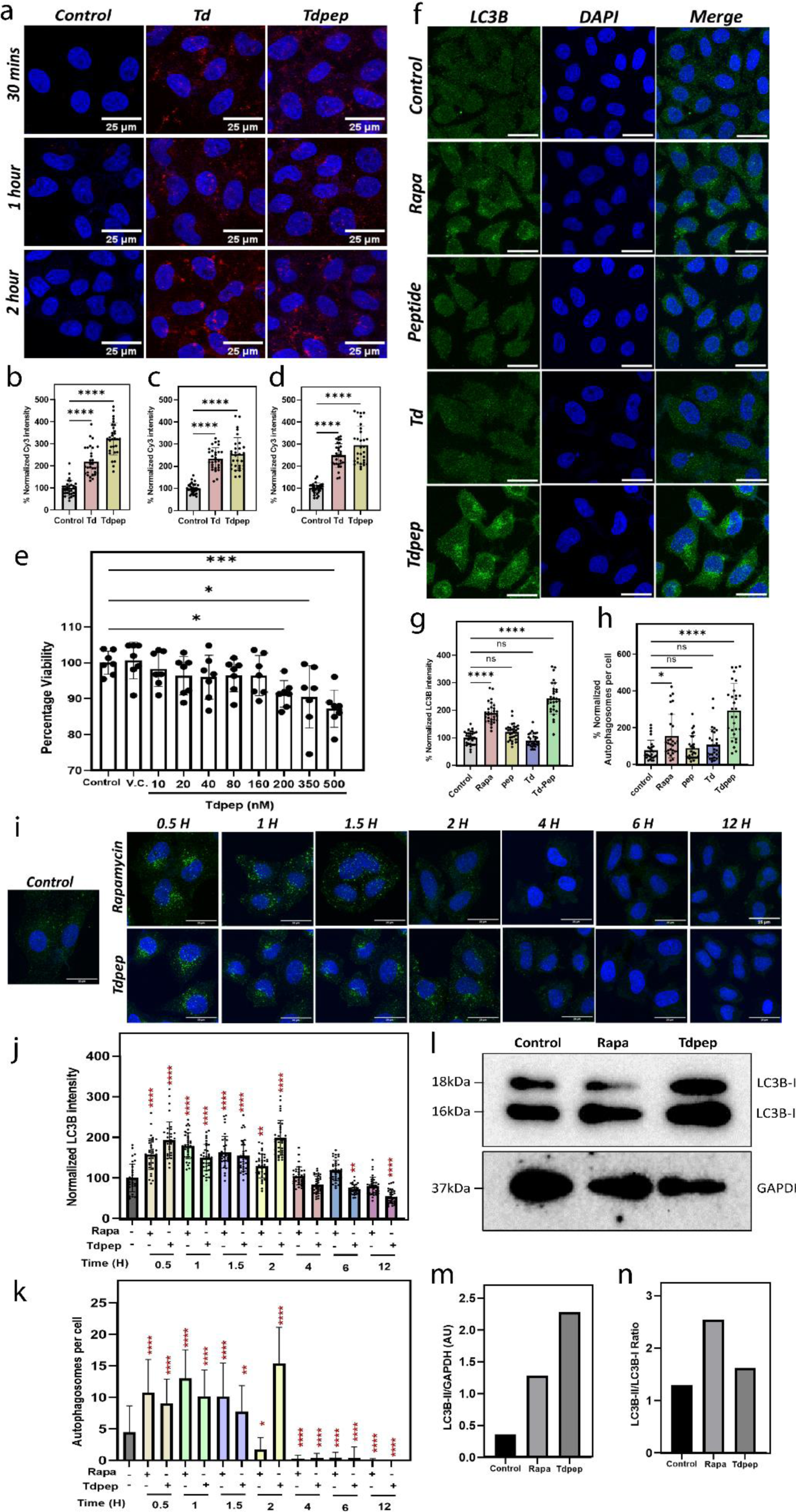
Tdpep nanosystem induces autophagy invitro. a. Confocal images of uptake of Td and Tdpep nanosystem in HeLa cells. The scale bar represents 25µm. b. Quantification of Td and Tdpep nanosystem uptake after 30 minutes. Histobars represents mean ± SD of 30 cells. c. Quantification of Td and Tdpep nanosystem uptake after 1 hour. Histobars represents mean ± SD of 30 cells. d. Quantification of Td and Tdpep nanosystem uptake after 2 hours. Histobars represents mean ± SD of 30 cells. e. MTT assay of Tdpep nanosystem in HeLa cells. V.C. represents vehicle control of 1X PBS used in 500nM of Tdpep. f. Confocal images of LC3B immunofluorescence staining. Scale bar represents 25µm. g. Quantification of LC3B intensity. Histobars represents mean ± SD of 30 cells. h. Quantification of autophagosomes per cell. Histobars represents mean ± SD of 30 cells. i. Confocal images of time-dependent autophagy induction analysis through LC3B immunofluorescence. The scale bar represents 25µm. j. Quantification of LC3B intensity across different timepoints after Tdpep treatment. Histobars represents mean ± SD of 30 cells. k. Quantification of autophagosomes per cell. Histobars represents mean ± SEM of 30 cells. l. LC3B lipidation assay with western blot. m. Quantification of LC3B-II band intensity normalized with respect to GAPDH band intensity. n. Quantifcation of LC3B-I to LC3B-II conversion ratio.

Before assessing autophagy induction, to ensure biocompatibility we evaluated the cytotoxicity of the Tdpep nanosystem via MTT assay. HeLa cells were treated with varying concentrations of the Tdpep system (10, 20, 40, 80, 160, 200, 350, and 500 nM) for 24 hours. Results showed a significant reduction in cellular viability beyond 200 nM. At 200 nM, over 90% of cells remained viable, establishing this concentration as optimal for further studies on autophagy induction (**Figure 3e**).

To investigate the autophagy-inducing potential of the Tdpep nanosystem, rapamycin (1 µM), a known potent autophagy inducer, was employed as a positive control. LC3B, a classical autophagosome marker, was used to monitor autophagy through immunofluorescence staining. We quantified total LC3B intensity and autophagosomes per cell, revealing that 200 nM Tdpep treatment led to a significant increase in autophagy induction compared to control, similar to the induction observed with 1 µM rapamycin. Importantly, neither the Td nanosystem nor the BH3 peptide alone induced significant autophagy, highlighting the specific role of the Tdpep conjugate in autophagy activation (**Figure 3f, 3g, 3h**).

Further investigation into the kinetics of autophagy induction by Tdpep was conducted using a time-dependent assay with Tdpep nanosystem. LC3B immunostaining showed that autophagy induction began at 30 minutes post-treatment with Tdpep, reaching a peak at 2 hours. Following this peak, autophagy levels returned to baseline, indicating a transient induction. This transient autophagic response was consistent with rapamycin treatment, where autophagy peaked at 90 minutes and then reverted to baseline levels (**Figure 3i, 3j, 3k**). The rapid yet temporary increase in autophagy is attributed to the limited stability of the Tdpep system, which maintained stability for approximately 2 hours (**Supplementary information, Figure 1**).

To further validate Tdpep-induced autophagy, an LC3B lipidation assay was performed. Upon autophagy induction, cytosolic LC3B-I is modified by conjugation with phosphatidylethanolamine (PE) to form membrane-bound LC3B-II, which localizes to autophagosomes. The LC3B-I to LC3B-II conversion ratio was assessed by western blotting, revealing an increased LC3B- II/LC3B-I ratio in cells treated with both Tdpep and rapamycin, indicative of enhanced LC3B lipidation and autophagy induction (**Figure 3l, 3m, 3n**). Collectively, these findings demonstrate that the Tdpep nanosystem effectively induces autophagy in vitro.

### 2.3 Tdpep nanosystem as an autophagy inducer and enhancer of autophagic flux

To determine whether the observed autophagosome accumulation following Tdpep treatment results from autophagy induction or from impaired autophagic flux, an autophagy flux assay was performed in the presence of the autophagy flux inhibitor Bafilomycin A1 (BafA1). BafA1, a specific inhibitor of the vacuolar-type H+-ATPase (V-ATPase) on lysosomes, prevents the fusion of autophagosomes with lysosomes, effectively blocking autophagic flux(24). This approach enabled us to differentiate between increased autophagosome formation and potential accumulation due to autophagy flux blockage (**Figure 4a**).

**Figure 4:**
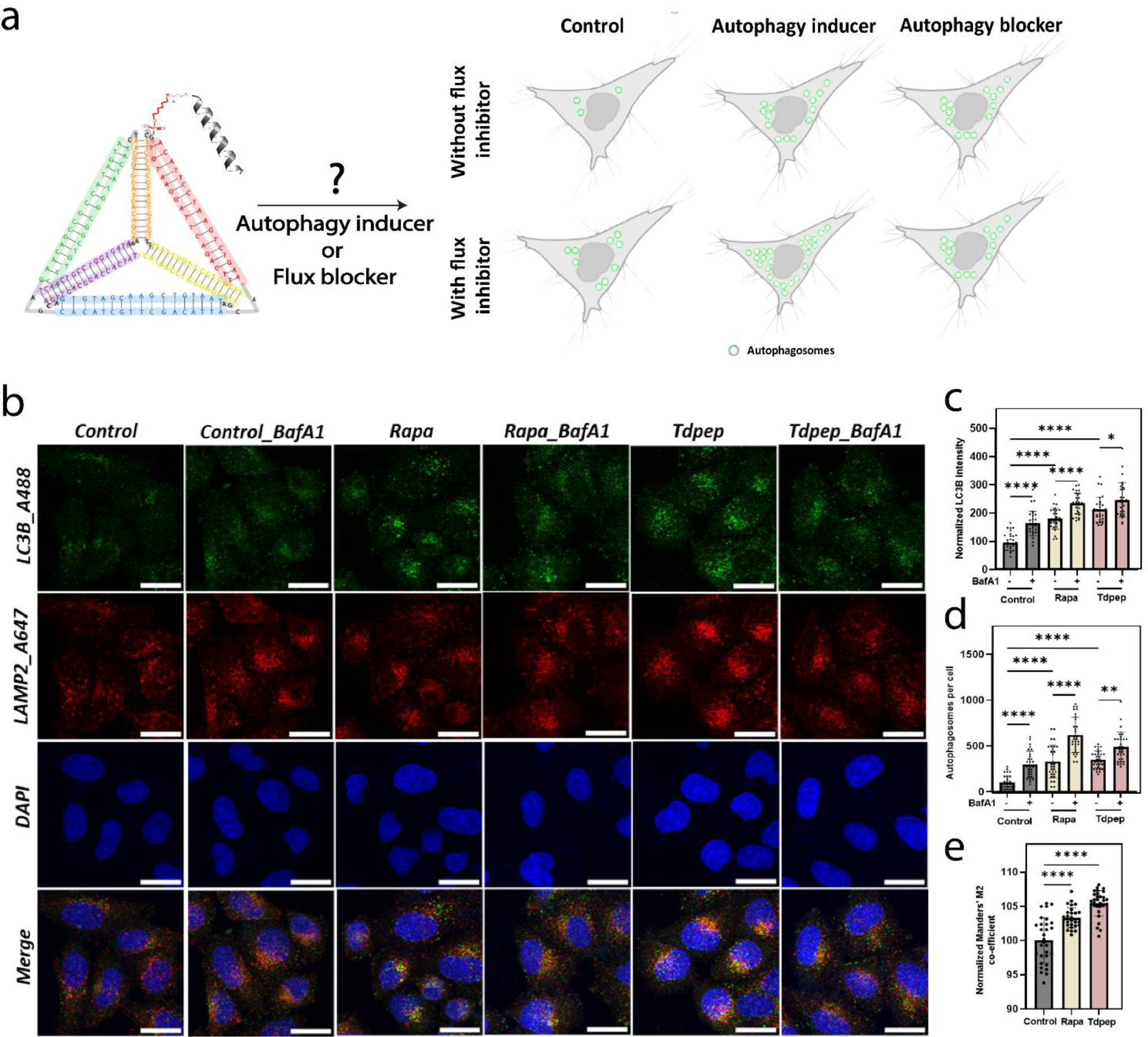
Autophagy flux analysis for Tdpep nanosystem: a. Schematics of the analyzing whether Tdpep nanosystem is an autophagy inducer or a blocker. b. Confocal images of the double immunofluorescence staining with LC3B and LAMP2. Scale bar represents 25µm. c. Quantification of LC3B intensity. Histobars represents mean ± SD of 30 cells. d. Quantification of autophagosomes per cell. Histobars represents mean ± SD of 30 cells. e. Quantification of the normalized M2 coefficient (percent of autophagosomes fused with lysosomes). Histobars represents mean ± SD of 30 cells.

Double immunofluorescence labeling of autophagosomes with LC3B and lysosomes with the lysosomal-associated membrane protein 2 (LAMP2) was performed to visualize autophagic flux. In control cells treated with BafA1, we observed marked accumulation of autophagosomes, as evidenced by increased LC3B puncta, confirming that BafA1 effectively inhibited autophagosome-lysosome fusion (**Figure 4b, 4c**). A corresponding decrease in the Mander’s M2 coefficient, which measures the percentage of autophagosomes co-localized with lysosomes, further confirmed the blockage of autophagosome-lysosome fusion upon BafA1 treatment (**Supplementary Information, Figure 2a, 2b, 2c**).

When autophagosome-lysosome fusion was inhibited with BafA1, cells treated with rapamycin (positive control) and Tdpep exhibited significantly higher levels of autophagosome accumulation compared to their respective treatments without BafA1 (**Figure 4b, 4c, 4d**). This increase in autophagosome accumulation upon BafA1 treatment indicates that both rapamycin and Tdpep stimulate autophagic activity, leading to higher autophagosome production.

To further quantify autophagic flux, we analyzed the Mander’s M2 coefficient in Tdpep- and rapamycin-treated cells. An increased M2 coefficient in Tdpep- and rapamycin-treated samples relative to control supports the conclusion that both treatments enhance autophagic flux by promoting autophagosome formation and subsequent autophagosome-lysosome fusion (**Figure 4b, 4e**).

In summary, these results demonstrate that Tdpep functions not only as an autophagy inducer but also as a robust enhancer of autophagic flux. The ability of Tdpep to enhance autophagic flux without causing flux blockage underscores its potential as an efficient autophagy inducer.

### 2.4 Selective induction of autophagy over apoptosis and ROS clearance by Tdpep nanosystem

The relationship between autophagy and apoptosis is complex, with both pathways often activated by overlapping stimuli and sharing common molecular regulators that enable cross-talk and influence cellular fate decisions (25). Given that many autophagy inducers can also activate apoptotic pathways, it is essential to confirm that a putative autophagy inducer, such as the Tdpep nanosystem, selectively stimulates autophagy without inducing apoptosis.

To investigate the specificity of Tdpep for autophagy induction over apoptosis, we performed flow cytometry analysis using Annexin V and propidium iodide (PI) staining. This assay enables distinction between early and late apoptotic populations: cells positive for Annexin V alone indicate early apoptosis, while cells positive for both Annexin V and PI mark late apoptotic stages. As a positive control, cells were treated at 55°C for 15 minutes to induce apoptosis. This treatment led to a significant increase in the Annexin V^+^/PI^+^ population, indicating robust late apoptosis induction, although early apoptotic cells were less prominently detected, likely due to the intensity of the thermal stress (**Figure 5a, 5b, 5c**).

**Figure 5.**
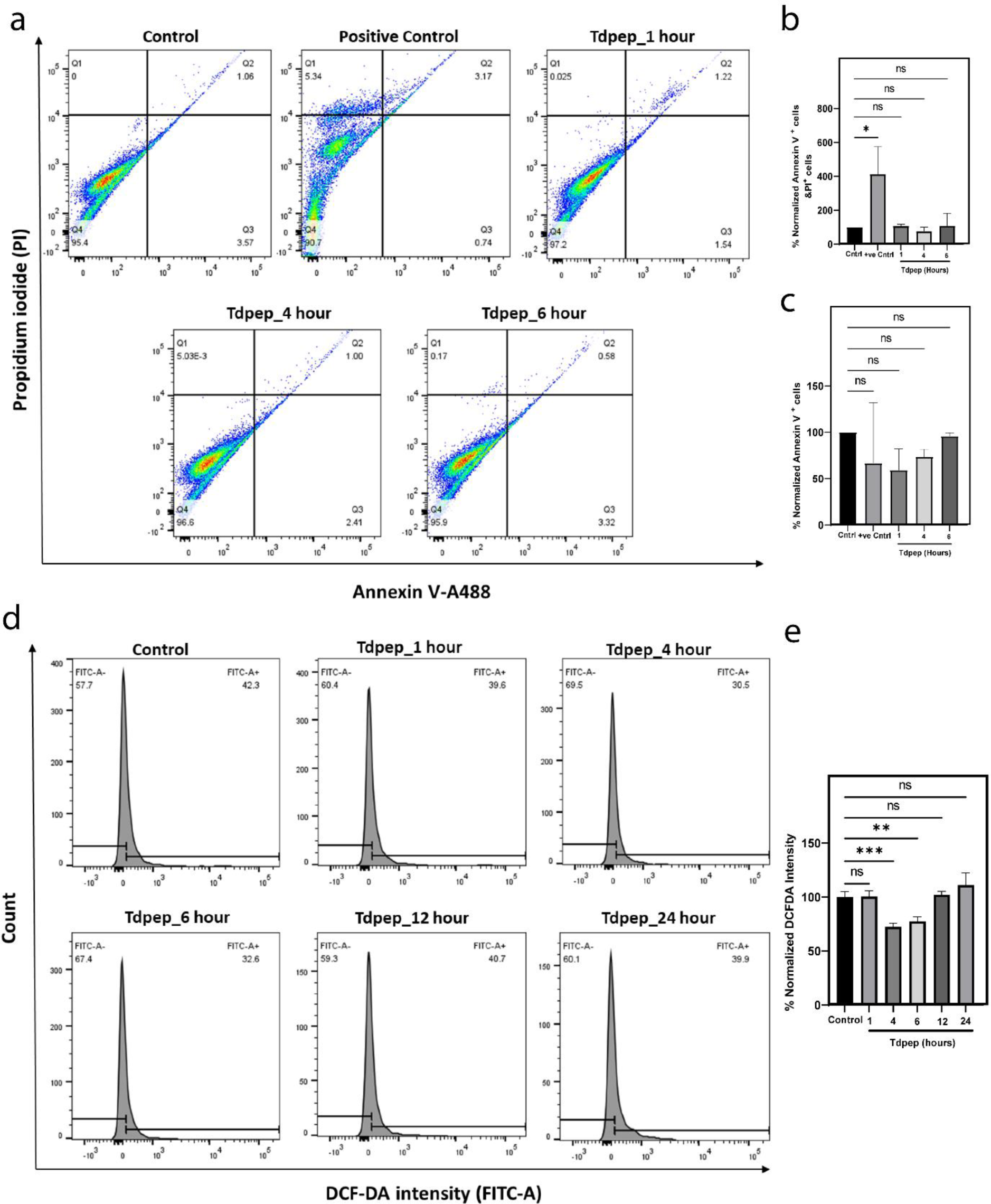
**Apoptosis and ROS level analysis**. a. Flow cytometry analysis of apoptosis of the gated population of single cells. b. Quantification of late apoptotic population. c. Quantification of early apoptotic population. 20,000 cells were analyzed per experiment. The experiment was carried out twice. d. Flow cytometry analysis of DCF-DA for the gated population of single cells. e. Quantification of DCF-DA intensity. 20,000 cells were analyzed per experiment. The experiment was carried out thrice. Histobars represents mean ± SEM.

We then assessed apoptosis induction in Tdpep-treated cells over a 6-hour period. Analysis revealed no significant increase in either early or late apoptotic populations, as indicated by the lack of detectable Annexin V^+^ or Annexin V^+^/PI^+^ cells in the Tdpep-treated samples (**Figure 5a, 5b, 5c**). These findings suggest that Tdpep does not stimulate apoptotic pathways and supports its role as a specific autophagy inducer without concurrent apoptosis activation.

To further substantiate the induction of autophagy by Tdpep, we evaluated the cellular outcome of autophagy activation by assessing intracellular reactive oxygen species (ROS) levels. ROS levels can provide insight into autophagy activity, as autophagic clearance of damaged mitochondria typically results in a reduction of cellular ROS (26). For this purpose, cellular ROS was quantified using 2′,7′-dichlorofluorescin diacetate (DCF-DA), which become fluorescent upon oxidation by ROS, and analyzed via flow cytometry and confocal microscopy.

Flow cytometry analysis of DCF-DA fluorescence showed a notable reduction in FITC^+^ cell population 4 hours after Tdpep treatment, signifying a decrease in cellular ROS levels (**Figure 5d, 5e**). This reduction in ROS was further corroborated by confocal microscopy, where a significant decline in DCF-DA fluorescence intensity was observed (**Supplementary information, Figure 3a, 3b**), consistent with the flow cytometry findings. These complementary results support that Tdpep treatment effectively reduces ROS levels, aligning with autophagy-mediated clearance mechanisms and reinforcing Tdpep’s role as a specific autophagy inducer.

### 2.5 Tdpep induces autophagy in vivo

To assess the in vivo autophagy-inducing potential of the Tdpep nanosystem, we utilized Danio rerio (zebrafish) larvae as a model organism. Monodansylcadaverine (MDC), a fluorescent probe that selectively labels acidic vesicles, including autophagosomes was employed to monitor autophagy induction (27). Following treatment with Tdpep or rapamycin (positive control), zebrafish larvae were examined for MDC staining (**Figure 6a**). Quantitative fluorescence analysis showed a significant increase in overall MDC fluorescence intensity in larvae treated with Tdpep compared to untreated controls, indicating a higher accumulation of acidic vesicles (**Figure 6b**). Furthermore, the mean MDC fluorescence intensity per unit area was elevated in the Tdpep and rapamycin-treated groups, supporting the Tdpep nanosystem’s efficacy in inducing autophagy in vivo (**Figure 6c**). These results substantiate Tdpep’s role in promoting autophagy.

**Figure 6.**
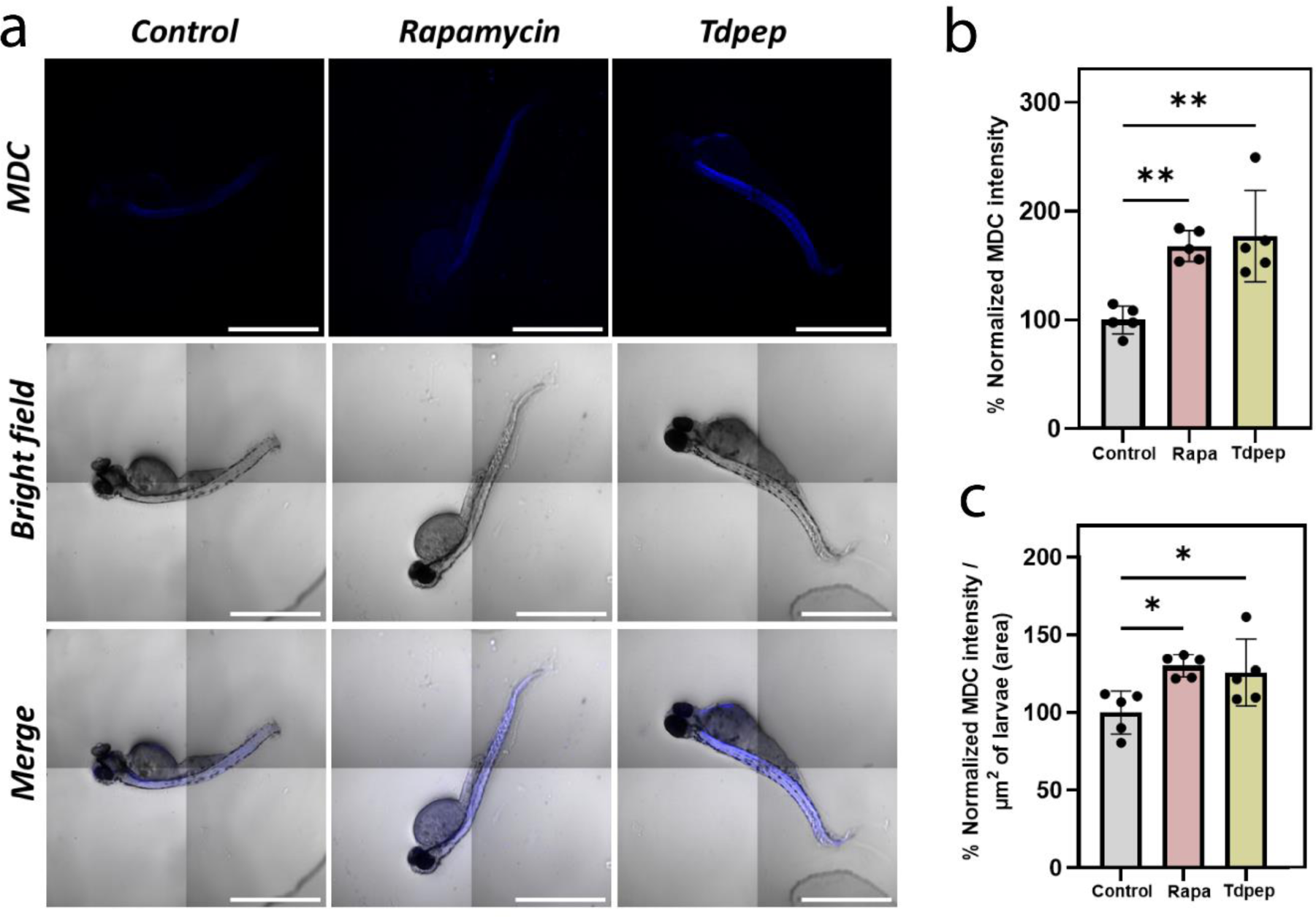
Tdpep nanosystem induces autophagy in vivo. a. Confocal images of MDC staining in the Danio rerio larvae. Scale bar represents 1000µm. b. Quantification of MDC intensity. Histobars represent mean ± SD for 5 Danio rerio larvae. c. Quantification of MDC intensity per unit area of the larvae. Histobars represent mean ± SD for 5 Danio rerio larvae.

## 3. Conclusion

Our study presents the DNA tetrahedron- peptide (Tdpep) nanosystem as a tool for autophagy induction via disruption of the Beclin 1-Bcl2 interaction, a central checkpoint in autophagy regulation. Unlike many conventional autophagy inducers that simultaneously activate apoptosis, the Tdpep nanosystem selectively induces autophagy without significant apoptotic cell death, as verified by Annexin V/PI assays. This specificity offers a key therapeutic advantage, especially in clinical settings where unintended apoptotic activation could compromise cell survival and overall therapeutic outcomes. Additionally, Tdpep treatment resulted in enhanced ROS clearance, a downstream effect of autophagy induction, thereby supporting its potential as a modulator of cellular homeostasis through effective autophagic flux (**Figure 7**).

**Figure 7:**
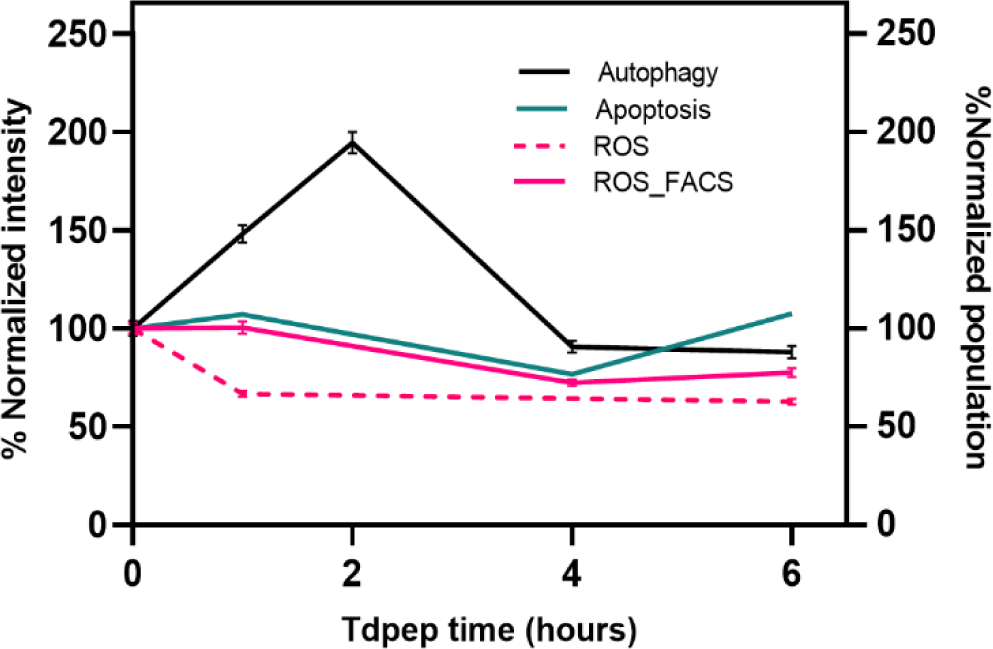
Summary of the study showing the dynamics of specific autophagy induction

In vivo studies in *Danio rerio* larvae further corroborated the efficacy of Tdpep, showing increased MDC staining in treated larvae, comparable to the well-known autophagy inducer rapamycin. This validation in a model organism highlights Tdpep’s functionality and efficacy in complex biological systems, confirming its relevance as a targeted autophagy inducer that could be adapted for use across different disease states.

An important aspect of Tdpep’s mechanism is the transient nature of autophagy induction, with autophagy peaking shortly after treatment and then subsiding. This transient effect could be advantageous for avoiding cellular stress and maintaining metabolic balance, as prolonged or continuous autophagy activation can strain cellular resources and lead to detrimental effects. However, in cases of chronic or progressive diseases where extended autophagy stimulation may enhance therapeutic benefits, enhancing Tdpep’s stability could be achieved by developing a D-peptide variant. D-peptides, due to their resistance to enzymatic degradation, would increase Tdpep’s half-life and extend the duration of autophagy induction, potentially broadening its application to diseases that require sustained autophagic clearance.

This research positions Tdpep as a promising candidate for therapeutic intervention in a range of diseases associated with autophagy dysregulation, including neurodegenerative disorders, cancers, and metabolic diseases. Future studies could focus on refining Tdpep’s structure for enhanced stability and optimizing delivery systems to target specific tissues or disease sites, ultimately paving the way for more precise and effective autophagy-based therapies. The DNA tetrahedron nanostructure, with its tunable design and functional versatility, combined with the selective autophagy-inducing peptide, provides a robust platform for further exploration into targeted autophagy modulation, potentially can revolutionize the treatment landscape for autophagy-related diseases.

## 4. Materials and methods

### 4.1 Materials and reagents

All the amino acids and reagents of peptide synthesis was purchased from Sigma Aldrich. All the oligonucleotide strands (M1, M2, M3, M4, M4Cy3) were purchased from Sigma. Dulbecco’s Modified Eagle Medium (DMEM), Fetal bovine serum (FBS), PenStrep, Trypsin-EDTA (0.25%), and collagen were purchased from Gibco. Nuclease free water, ammonium persulfate, ethidium bromide, TEMED, Triton-X, paraformaldehyde, and adherent cell culture dishes were purchased from Himedia. Tris-Acetate EDTA (TAE), Acrylamide/bis(acrylamide) sol 40% were purchased from GeNei. Magnesium chloride was ordered from SRL, India. SPDP (N-succinimidyl 3-(2- pyridyldithio) propionate) was purchased from invitrogen. Bafilomycin A1 was purchased from Sigma Aldrich. Rapamycin was purchased from TCI.

### 4.2 Structure prediction and Docking

The tertiary structure of the peptide was predicted with AlphaFold2 ColabFold v1.5.5. The peptide sequence was inputted as query, and the structure was predicted with template mode pdb100, keeping all other parameters default. The generated pdb files were visualized using UCSF Chimera. For docking analysis, the x-ray crystallography based derived structure of Bcl2 (5VAU) was derived from PDB database. It was docked with the predicted peptide structure using Haddock 2.4 online server by providing the ambiguous restrains. (**Supplementary information, Table 1**). All other parameters were kept default for docking analysis.

### 4.3 Synthesis of peptide

The 21 amino acids long peptide CTMENLSRRLKVTGDLFDIMS was synthesized in-house with Fmoc-based solid phase peptide synthesis. The peptide was synthesized on Fmoc-protected rink amide resin with a loading capacity of 0.59mmol/g. The reaction scale was 100 mg. The resin was weighed and swelled in anhydrous DCM for 3 hours, followed by swelling in DMF for 1 hour. After draining, the resin was washed with DCM and DMF alternatively. The resin was subjected to Fmoc removal and coupling of the first amino acid through the following steps:

1. Decoupling of Fmoc: The Fmoc was decoupled with the 5% piperazine and DBU in DMF. Decoupling was carried out for three cycles with incubation times of 12, 10, and 5 minutes, respectively. After each decoupling, DCM-DMF washes were carried out.
2. Coupling: The Fmoc-protected amino acid coupling was carried out by the DIC/oxyma method. The amino acid and oxyma were taken in 3 equivalents dissolved in DMF and preactivated with 3 equivalents of DIC for 5 minutes with mild shaking. Double coupling was carried out to attach the activated amino acid to the growing peptidyl resin. Each coupling was carried out for 1 hour. After coupling, the peptidyl resin was given DCM-DMF washes.

Steps 1 & 2 were repeated until the desired peptide sequence was obtained. Post-synthesis, the peptidyl resin was washed with methanol. The peptide was cleaved from the resin with the cleavage cocktail consisting of 92.5% TFA, 2.5% DODT, 2.5% TIPS, and 2.5% water for 3 hours with continuous vortexing. The resin was removed with filtration, and the filtrate was evaporated under nitrogen flow nearly to 1/5^th^ of the total volume. The peptides were precipitated with the ice-cold diethyl ether and centrifuged at 8000rpm for 30 minutes at 4°C. The supernatant was decanted, and the pellet was dissolved in 50% acetonitrile in water and further frozen and lyophilized. The lyophilized powder was stored at -20°C for further use. The peptides were characterized with MALDI-ToF.

### 4.4 Coupling of peptide and DNA oligo

The cysteine-modified peptide and 5’ amino-modified M1 oligo were coupled using the SPDP cross-linking. Briefly, the 20µM amino-modified M1 strand was incubated with 2mM of SPDP in PBS-EDTA with continuous mixing for 1 hour at room temperature. The unreacted SPDP was desalted with a 7K Zeba spin desalting column. The peptide was dissolved in PBS-EDTA. Further, M1-SPDP is incubated with 5 equivalents of peptide in an equimolar ratio to get the final working concentration of 10µM of M1 coupled to the peptide. The unreacted peptide was removed through desalting.

### 4.5 Synthesis of DNA tetrahedron coupled peptide

Four oligo strands (M1, M2, M3, M4) were used to synthesize DNA tetrahedron (**supplementary information, table 3**). The oligo strands were reconstituted with nuclease-free water to 100µM at 70°C for 1 hour 30 minutes. The M1 strand was coupled with peptide. The other three strands were diluted to 10µM working concentration. DNA tetrahedron (Td) was formed through one-pot synthesis, where all four oligo strands (M1-pep, M2, M3, M4) were added in the equimolar ratio with 2 mM MgCl2. The reaction was performed at 95°C for 10 minutes and immediately cooled to 4°C. The final concentration of Td was 2.5µM and was stored at 4°C until further use.

### 4.6 MALDI-TOF

For MALDI-TOF mass spectrometry analysis, α-Cyano-4-hydroxycinnamic acid (HCCA) was used as the matrix. The synthesized peptide solution was mixed with the HCCA matrix solution in a 1:1 volume ratio, and 1 µL of this mixture was applied to a MALDI target plate. The sample spot was air-dried at room temperature to form a homogeneous crystalline layer. MALDI-TOF analysis was conducted in positive ion mode with laser desorption/ionization, optimized for peptide detection, and spectra were acquired.

### 4.7 Electrophoretic Mobility Shift Assay

The Electrophoretic Mobility Shift Assay (EMSA) was used to estimate the formation of higher-order structures. 10% Native-PAGE was used to carry out EMSA. 5µl of the DNA Tdpep samples were loaded along with 3µl of loading buffer and 1.5µl of 6X loading dye. The Native gel was run with a constant voltage of 70V for 90 minutes. The gel was stained with 0.32µg/ml of ethidium bromide for 10 minutes and visualized using the Gel documentation system (Biorad ChemiDoc MP Imaging System).

### 4.8 UV-Visible Spectroscopy

UV-Visible (UV-VIS) spectroscopy was used for characterizing the coupling of M1 and M1-pep, assessing the wavelength peak at 260nm (DNA) and 220nm (peptide). Samples were diluted at a 1:10 ratio, and 1ml was analyzed using Test Right UV-Visible spectroscopy. Measurements were taken in triplicate for accuracy.

### 4.9 Dynamic Light Scattering

Dynamic light scattering (DLS) was used for size-based characterization of Td and Tdpep, assessing the hydrodynamic size of the structures. Samples were diluted at a 1:10 ratio, and 1ml was analyzed using a Malvern Zetasizer Nano ZS. Measurements were taken in triplicate for accuracy.

### 4.10 Atomic Force Microscopy

All samples were diluted 10X in nuclease-free water. The diluted samples were subsequently vacuum-dried on the mica sheets glued to a glass slide for 1 day. After drying, the samples were analyzed using an Atomic Force Microscope (Bruker) in air-tapping mode using a cantilever tip (Force Mod, 1 nm Tip radius, 80 KHz). Before analysis, the cantilever tip was pre-calibrated, and the phase difference was set to zero. The scanning line rate of 1 Hz and a 30% set point were used for image acquisition during analysis. JPK data processing software was used for data analysis.

### 4.11 Stability

The stability of the Tdpep system was assessed with the serum stability assay. The Tdpep was incubated with 10% FBS at 37°C for different time points (0.5, 1, 2, 3, 4, 6 and 8 hours) and the reaction was immediately stopped by storing at -20°C. Then stability of samples was checked with 8% Native PAGE. 5µl of the DNA Tdpep samples were loaded along with 3µl of loading buffer and 1.5µl of 6X loading dye. The Native gel was run with a constant voltage of 75V for 60 minutes. The gel was stained with 0.32µg/ml of ethidium bromide for 10 minutes and visualized using the Gel documentation system (Biorad ChemiDoc MP Imaging System).

### 4.12 Cell culture

HeLa (Cervical cancer) cells were maintained in Dulbecco-modified eagle media (DMEM) media supplemented with 10% fetal bovine serum and 1% penicillin-streptomycin. The cells were incubated at 37°C in the humidified chamber with 5% CO2. Generally, the cells were passaged once 85-95% confluency was reached, and the media was changed as required.

### 4.13 Uptake studies

For the cellular uptake of the Tdpep study, the cells were seeded with the seeding density of 0.1 x 10^6^ cells per well onto 12mm coverslips and incubated for 24 hours. The cells were washed with 1X PBS, and the cells were incubated with 200nM of Td and Tdpep for different time points (30 minutes, 1 hour, and 2 hours) in serum-free media at 37°C. The incubated cells were washed with 1X PBS thrice to remove the excess and surface-bound Td and Tdpep nanostructures. Then, the cells were fixed with 4% paraformaldehyde at 37°C for 15 minutes. Further, the cells were washed with 1X PBS thrice, and the coverslips with cells were mounted on the glass slide with mowiol containing DAPI to mark the nucleus.

### 4.14 Cell viability

To assess cell viability, 20,000 cells were seeded per well and incubated at 37°C for 24 hours. The media was removed, and the cells were exposed to different concentrations of peptide and Tdpep for 24 hours at 37°C. After incubation, 0.5mg/ml of MTT ((3 (4,5-Dimethylthiazol-2-yl)- 2,5-diphenyl tetrazolium bromide) solution was added to each well and incubated at 37°C for 4 hours. After incubation, the MTT solution was removed, replaced with 100µl of DMSO, and incubated in the dark for 15 minutes to dissolve the formazan crystals. The absorbance at 562nm was measured with the plate reader (Byonoy). The percentage of cell viability is calculated as

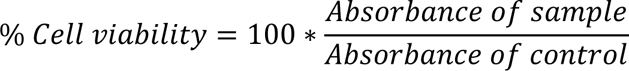

### 4.15 Immunofluorescence

For all the immunofluorescence studies, the cells were seeded onto the 12mm coverslips with the seeding density of 0.1 x 10^6^ cells and incubated at 37°C for 24 hours. After the cell attachment, the cells were treated with the respective treatment, and were fixed with 4% paraformaldehyde at 37°C for 15 minutes. Then the cells were permeabilized with 0.1% triton x- 100 for 15 minutes at 37°C. The cells were incubated with the blocking buffer (10%FBS, 0.05% triton x-100 in 1X PBS) for 1 hour at 37°C. The cells were incubated with the primary antibody at 1:100 for LC3B (CST, 2775s) and 1:200 for LAMP2 in blocking buffer for 2 hours at 37°C. The cells were washed with 1X PBS thrice. Further, the cells were incubated with the secondary antibody (Anti-rabbit A488 (invitogen, A11008), Anti-rabbit A647 (invitogen A21244), Anti-mouse A647 (CST, 4410s)) at a dilution of 1:1000 at room temperature for 2 hours. The cells were washed with 1X PBS thrice and the coverslips with the cells were mounted on the glass slide with mowiol and DAPI.

### 4.16 Western blot analysis

The cells were cultured in the 100mm petri dish. The cells were washed with ice-cold PBS and then scraped out in ice-cold PBS. The scrapped cells were pelleted and lysed with lysis buffer () supplemented with the protease inhibitor cocktail. The cells were incubated on ice for 30 minutes and then centrifuged at 9000rpm for 20 minutes at 4°C. The protein concentration of the supernatant was quantified using the Bradford assay. Then, the lysates were denatured at 95°C for 5 minutes. For western blot analysis, the samples were subjected to separation with 15% SDS-PAGE separation (5% stacking) with a constant voltage of 100V for 90 minutes. The resolved samples were transferred to the PVDF membrane with a constant ampere of 200mA for 55 minutes. Then, the PVDF membranes were blocked with 5% skimmed milk in TBST for 1 hour at room temperature. Then, the membrane was incubated with the primary antibody at the dilution of 1:1000 for LC3B (Invitrogen, MA5-42722) and 1:2000 for GAPDH (Invitrogen, MA5- 15738) overnight at 4°C with gentle shaking. Then, the membrane was washed thoroughly with TBST and further incubated with a HRP conjugated secondary antibody at a dilution of 1: 10,000 [(Anti-rabbit, Invitogen, 31460), Anti-mouse, Invitrogen, PA1-28568)] for 2 hours at room temperature. The membrane was washed with 1X TBST thrice, 15 minutes each, with intense shaking. The membrane was developed with an ECL chemiluminescence detection kit and visualized with the Gel documentation system (Biorad ChemiDoc MP Imaging System). The protein level on the blot was quantified using ImageJ software.

### 4.17 Apoptosis analysis

For the apoptosis analysis, the cells were seeded with the seeding density of 0.1 x 10^6^ cells per well and incubated for 24 hours. The cells were harvested with 0.25% trypsin and pelleted down. The apoptosis analysis was carried out with the dead cell apoptosis kits with Annexin V (V13241). The pelleted cells were dispersed in 1X annexin binding buffer with Annexin V and PI according to the kit protocol. Immediately the cells were analyzed for Annexin V and PI staining with flow cytometry.

### 4.18 Intracellular ROS measurement

The intracellular ROS was measured with DCF-DA. For flow cytometry analysis of ROS with DCF-DA, the cells were treated with 200nM of Tdpep for various time points (1, 4, 6, 12, and 24 hours). Following the treatment, the cells were washed with 1X PBS and incubated with 20µM of DCA-DA for 15 minutes. The cells were washed with 1X PBS and removed with 0.25% trypsin, pelleted down and dissolved in 1X PBS for further analysis with flow cytometry. For confocal analysis of ROS with DCF-DA, the cells were seeded with the seeding density of 0.1 x 10^6^ cells per well onto 12mm coverslips and incubated for 24 hours. The seeded cells were treated with 200nM of Tdpep and 1µM of rapamycin for 1 and 6 hours. Following the treatment, the cells were washed with 1X PBS and incubated with 10µM of DCA-DA for 30 minutes. The cells were washed with 1X PBS and fixed with 4% paraformaldehyde. The fixed cells were washed with 1X PBS and coverslips containing the cells were mounted on the slides with mowiol and DAPI.

### 4.19 *Danio rerio* (Zebrafish) husbandry and maintenance

The zebrafish (Assam wild-type) were reared from embryos to adults under controlled laboratory conditions, following ZFIN guidelines. A 14 h light/10 h dark cycle was maintained at 26–28 °C. Fish were housed in aerated 20 L tanks with water conditions optimized to mimic their natural habitat: pH 7–7.4, conductivity 250–350 μS, TDS 220–320 mg L⁻¹, salinity 210–310 mg L⁻¹, and dissolved oxygen > 6 mg L⁻¹, monitored via a PCD 650 multi-parameter instrument. The diet consisted of live Artemia (twice daily) and Aquafin basic flakes (once daily). For breeding, a 3:2 female-to-male ratio was used, and embryos were collected in E3 medium, then incubated at 28.0 °C.

### 4.20 Monodansyl cadaverine staining for autophagy assessment in *Danio rerio* larvae

After 72 hours post fertilization, the healthy larvae were selected for the experiment. The larvae were treated with 300nM Tdpep and 1µM rapamycin for 6 hours. Post treatment the larvae were washed with water. Then the larvae were treated with 500µM of monodansyl cadaverine for 30 minutes and washed with water. The larvae were fixed with 4% paraformaldehyde and washed thoroughly with water. Following it, the larvae were mounted onto the slides with mowiol.

### 4.21 Confocal microscopy imaging

All the fixed cells confocal imaging was carried out in the Leica TCS SP8 confocal laser scanning microscopy. The images were captured with the 63X oil immersion objective. The pinhole was kept constant as 1 Airy unit. The bit depth, laser power and detector gain was kept constant across different experimental condition in a particular study. All the images were taken at the resolution of 512 x 512. The DAPI, monodansyl cadaverine was excited at 405nm with Diode 405 laser, DCFDA and Alexa488 fluorophores was excited at 488nm with Argon laser, Cy3 was excited at 561nm with DPSS laser, and Alexa647 was excited at 633nm with He-Ne laser. The detector emission collection window was set corresponding to the emission spectra of the fluorophores. Z-stack imaging was carried out and the number of z-steps were set to be system optimized. In case of multiple fluorophore imaging, sequential scanning was employed.

### 4.22 Image processing and statistical analysis

The image analysis was done Fiji ImageJ software. the z-stacks were merged into 2D images using the z-projection method, specifically employing the maxi mum intensity projection type. Background correction was executed through the subtraction of the average background intensity. Following this correction, integrated density and raw integrated density were quantified.

For studying autophagy, an object-based methodology was employed to track autophagosomes. An auto threshold value was used across all images to identify autophagosomes and differentiate them from the image background. Furthermore, a watershed algorithm was employed to segment autophagosomes located in close proximity for accurate quantification. To estimate autophagosomes per cell, a region of interest was drawn by tracing the cell’s perimeter to restrict autophagosome count to single cell. The autophagosome within the ROI were quantified by analyzing the particles with the following criteria: circularity cutoff of 0.5–1 and particle size range of 0.5–1.8 µm^2^.

For co-localization analysis, the Mander’s co-efficient was analyzed as a quantifier of spatial overlap between two fluorophores A488 (autophagosomes-channel 2) and A647 (lysosomes- channel1). The 3D z-stack of the fluorophore channels were given as input in coloc2 plugin of ImageJ. The M2 co-efficient that estimated the fraction of autophagosomes co-localized/ co-occurred with the lysosome within the resolution of the single voxel was used for autophagosome-lysosome fusion analysis.

Data normalization and statistical analysis were performed using GraphPad Prism (Version 9.0). The data normalization process involved assigning the value zero as 0% across all datasets. Concurrently, the average integrated density value of the control group was established as 100%. The data were analyzed using unpaired, 2-tailed t test when comparing two experimental groups or analysis of variance (ANOVA) followed by Dunnett’s test when comparing more than two experimental groups. All the test was carried out under the assumption of normality.

## 5. Conflicts of interest

The authors declare no conflicts of interest.

## 6. Author contributions

HNA and DB conceptualized the study. DB acquired the funding. HNA carried out the investigation. NS carried out AFM imaging and quantification. KK carried out the studies in in vivo model. HNA analyzed the data. AK provided the facilities and supervised the in vivo experiments. SG provided the facilities and supervised in the peptide synthesis. DB supervised the entire project. HNA wrote the original manuscript, which was reviewed and edited by all the authors.

## Acknowledgement

We sincerely thank all the members of the DB group for critically reading the manuscript and their valuable feedback. HN thank the Ministry of Education, GoI for the PMRF fellowship. KK thank DST Nanomission postdoctoral fellowship. NS acknowledge the MHRD for the PhD fellowship. DB thank SERB-CRG, MoES-STARS, Gujcost-DST, GSBTM and IITGN for the research grants. We sincerely thank Dr. Sharad Gupta for providing the peptide synthesis facility and Dr. Ashutosh kumar for the zebra fish facility. IIT Gandhinagar CIF facilities are duly acknowledged for Confocal, AFM, DLS MALDI, and FACS facilities.

## Notes

### Competing Interest Statement

The authors have declared no competing interest.

## References

1. Mizushima N, Komatsu M. Autophagy: Renovation of Cells and Tissues. Cell. 2011 Nov 11;147(4):728–41.

2. Li W, He P, Huang Y, Li YF, Lu J, Li M, et al. Selective autophagy of intracellular organelles: recent research advances. Theranostics. 2021 Jan 1;11(1):222.

3. Lamark T, Johansen T. Aggrephagy: Selective Disposal of Protein Aggregates by Macroautophagy. International Journal of Cell Biology. 2012 Mar 22;2012:736905.

4. Autophagy during viral infection — a double-edged sword | Nature Reviews Microbiology [Internet]. [cited 2024 Nov 1]. Available from: https://www.nature.com/articles/s41579-018-0003-6

5. Chen T, Tu S, Ding L, Jin M, Chen H, Zhou H. The role of autophagy in viral infections. Journal of Biomedical Science. 2023 Jan 18;30(1):5.

6. Bacteria–autophagy interplay: a battle for survival | Nature Reviews Microbiology [Internet]. [cited 2024 Nov 1]. Available from: https://www.nature.com/articles/nrmicro3160

7. Levine B, Packer M, Codogno P. Development of autophagy inducers in clinical medicine. J Clin Invest. 2015 Jan 2;125(1):14–24.

8. Zhang Z, Yang X, Song YQ, Tu J. Autophagy in Alzheimer’s disease pathogenesis: Therapeutic potential and future perspectives. Ageing Research Reviews. 2021 Dec 1;72:101464.

9. Lynch-Day MA, Mao K, Wang K, Zhao M, Klionsky DJ. The Role of Autophagy in Parkinson’s Disease. Cold Spring Harbor Perspectives in Medicine. 2012 Apr;2(4):a009357.

10. Jiang B, Zhou X, Yang T, Wang L, Feng L, Wang Z, et al. The role of autophagy in cardiovascular disease: Cross-interference of signaling pathways and underlying therapeutic targets. Frontiers in Cardiovascular Medicine. 2023 Mar 29;10:1088575.

11. Xu J, Kitada M, Ogura Y, Koya D. Relationship Between Autophagy and Metabolic Syndrome Characteristics in the Pathogenesis of Atherosclerosis. Frontiers in Cell and Developmental Biology. 2021 Apr 15;9:641852.

12. Autophagy in metabolic syndrome: breaking the wheel by targeting the renin– angiotensin system | Cell Death & Disease [Internet]. [cited 2024 Nov 1]. Available from: https://www.nature.com/articles/s41419-020-2275-9

13. Frontiers | The Double-Edge Sword of Autophagy in Cancer: From Tumor Suppression to Pro-tumor Activity [Internet]. [cited 2024 Nov 1]. Available from: https://www.frontiersin.org/journals/oncology/articles/10.3389/fonc.2020.578418/full

14. Ahmadi-Dehlaghi F, Mohammadi P, Valipour E, Pournaghi P, Kiani S, Mansouri K. Autophagy: A challengeable paradox in cancer treatment. Cancer Medicine. 2023;12(10):11542–69.

15. Bhutia SK, Mukhopadhyay S, Sinha N, Das DN, Panda PK, Patra SK, et al. Autophagy: Cancer’s Friend or Foe? Advances in cancer research. 2013;118:61.

16. Galluzzi L, Pedro JMBS, Levine B, Green DR, Kroemer G. Pharmacological modulation of autophagy: therapeutic potential and persisting obstacles. Nature reviews Drug discovery. 2017 May 19;16(7):487.

17. Florance I, Cordani M, Pashootan P, Moosavi MA, Zarrabi A, Chandrasekaran N. The impact of nanomaterials on autophagy across health and disease conditions. Cell Mol Life Sci. 2024 Apr 17;81(1):184.

18. Li Y, Ju D. The Role of Autophagy in Nanoparticles-Induced Toxicity and Its Related Cellular and Molecular Mechanisms. Adv Exp Med Biol. 2018;1048:71–84.

19. Paskeh MDA, Entezari M, Clark C, Zabolian A, Ranjbar E, Farahani MV, et al. Targeted regulation of autophagy using nanoparticles: New insight into cancer therapy. Biochimica et Biophysica Acta (BBA) - Molecular Basis of Disease. 2022 Mar 1;1868(3):166326.

20. Fernández ÁF, Sebti S, Wei Y, Zou Z, Shi M, McMillan KL, et al. Disruption of the beclin 1-BCL2 autophagy regulatory complex promotes longevity in mice. Nature. 2018 Jun;558(7708):136–40.

21. Dahle L, Vaswani P, Bhatia D. Tetrahedral DNA nanocages as delivery agent for biological and biomedical applications. Nano and Medical Materials. 2023 Nov 22;3(2):151– 151.

22. Sharma A, Vaswani P, Bhatia D. Revolutionizing cancer therapy using tetrahedral DNA nanostructures as intelligent drug delivery systems. Nanoscale Advances. 2024 May 28;6(15):3714.

23. Wang W, Lin M, Wang W, Shen Z, Wu ZS. DNA tetrahedral nanostructures for the biomedical application and spatial orientation of biomolecules. Bioactive Materials. 2024 Mar 1;33:279–310.

24. Mauvezin C, Neufeld TP. Bafilomycin A1 disrupts autophagic flux by inhibiting both V- ATPase-dependent acidification and Ca-P60A/SERCA-dependent autophagosome-lysosome fusion. Autophagy. 2015 Sep 14;11(8):1437.

25. Sorice M. Crosstalk of Autophagy and Apoptosis. Cells. 2022 Apr 28;11(9):1479.

26. Redza-Dutordoir M, Averill-Bates DA. Interactions between reactive oxygen species and autophagy: Special issue: Death mechanisms in cellular homeostasis. Biochimica et Biophysica Acta (BBA) - Molecular Cell Research. 2021 Jul 1;1868(8):119041.

27. Biederbick A, Kern HF, Elsässer HP. Monodansylcadaverine (MDC) is a specific in vivo marker for autophagic vacuoles. Eur J Cell Biol. 1995 Jan;66(1):3–14.

