## Supporting information for "Peptide modified, programmable DNA tetrahedra to modulate autophagy in biological systems"

**Engineering DNA tetrahedron peptide nanostructure to induce bulk autophagy invitro and invivo**

**Supplementary information**

**Table 1:** Restrains provided during molecular docking

| **Restrain type** | **Residue number**  **Bcl2** | **Residue number**  **BH3 peptide** | **Source** |
| --- | --- | --- | --- |
| Ambiguous | 104 | 9 | (1) |
|  | 202 | 14 |  |
|  | 112 | - |  |
|  | 146 | - |  |
|  | 108 | - |  |
|  | 107 | - |  |
|  | 136 | - |  |
|  | 140 | - |  |
|  | 143 | - |  |

**Table 2:** Predicted clusters for the docking model, their respective Haddock score, cluster size, RMSD values. The highlighted cluster was taken for further docking analysis

| **Docking** | **Cluster number** | **Cluster Size** | **Haddock Score** | **RMSD** |
| --- | --- | --- | --- | --- |
| BH3 peptide-Bcl2 | **1** | **44** | **-131.4 ± 9.2** | **0.4 ± 0.3** |
|  | 2 | 26 | -98.4 ± 6.7 | 2.2 ± 0.2 |
|  | 3 | 19 | -91.9 ± 5.8 | 6.5 ± 0.2 |
|  | 4 | 19 | -78.4 ±1.4 | 2.7 ± 0.5 |
|  | 5 | 13 | -80.4 ± 5.2 | 2.4 ± 0.3 |
|  | 7 | 7 | -86.9 ± 8.8 | 7.2 ± 0.1 |
|  | 8 | 6 | -92.0 ± 8.6 | 1.9 ± 0.5 |
|  | 9 | 6 | -78.0 ± 5.9 | 6.2 ± 0.0 |
|  | 10 | 1 | -85.5 ± 16.5 | 2.5 ± 0.2 |
|  | 11 | 5 | -71.0 ± 3.9 | 6.2 ± 0.1 |

**Table 3:** List of sequence of oligonucleotides for the DNA tetrahedron formation

| **Sequence ID** | **Sequence (5’-3’)** |
| --- | --- |
| M1 - No modification | ACATTCCTAAGTCTGAAACATTACAGCTTGCTACACGAGAAGAGCCGCCATAGTA |
| M1 - Amino modification | AmC6-ACATTCCTAAGTCTGAAACATTACAGCTTGCTACACGAGAAGAGCCGCCATAGTA |
| M2 | TATCACCAGGCAGTTGACAGTGTAGCAAGCTGTAATAGATGCGAGGGTCCAATAC |
| M3 | TCAACTGCCTGGTGATAAAACGACACTACGTGGGAATCTACTATGGCGGCTCTTC |
| M4 | TTCAGACTTAGGAATGTGCTTCCCACGTAGTGTCGTTTGTATTGGACCCTCGCAT |
| M4-Cy3 | Cy3-TTCAGACTTAGGAATGTGCTTCCCACGTAGTGTCGTTTGTATTGGACCCTCGCAT |

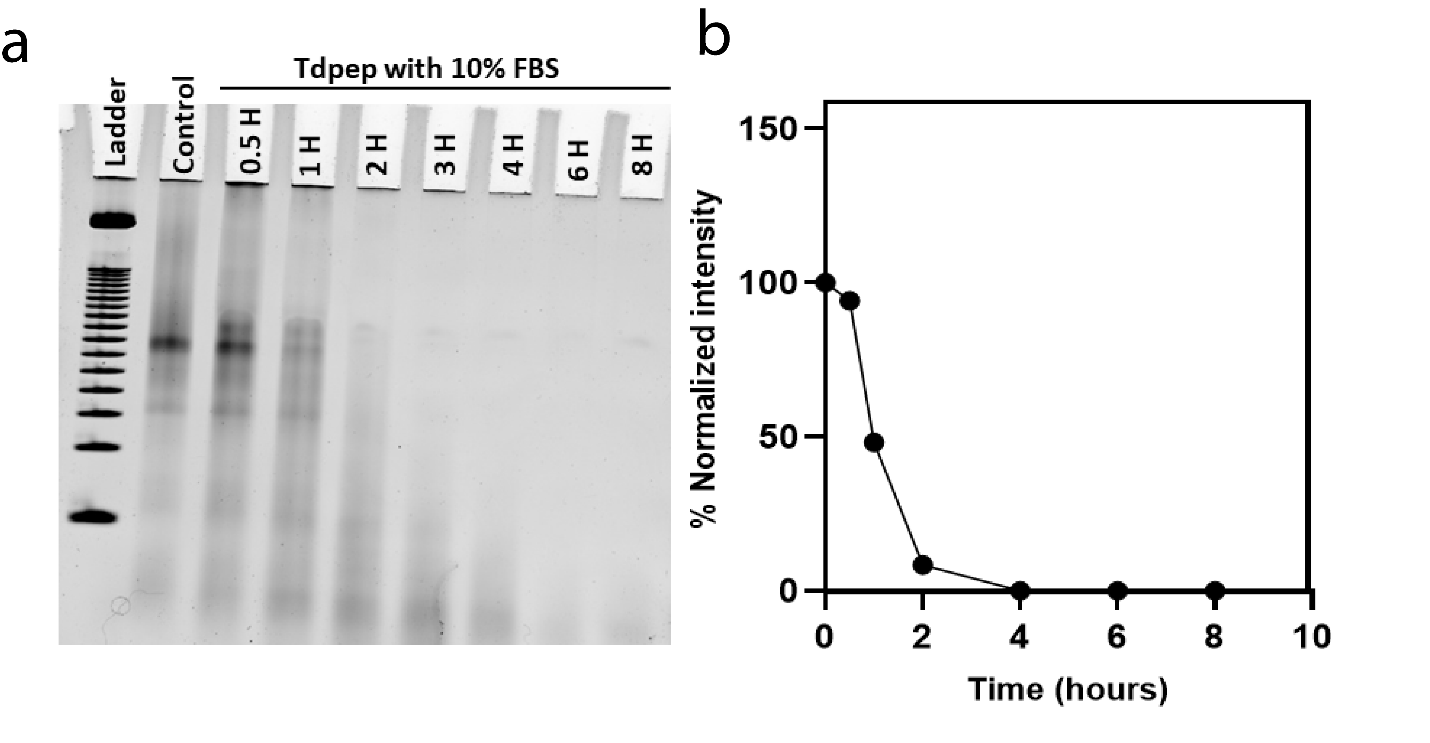

**Figure 1:** a. Serum stability assay in the 8% Native PAGE. b. Quantification of the band intensities of the serum stability

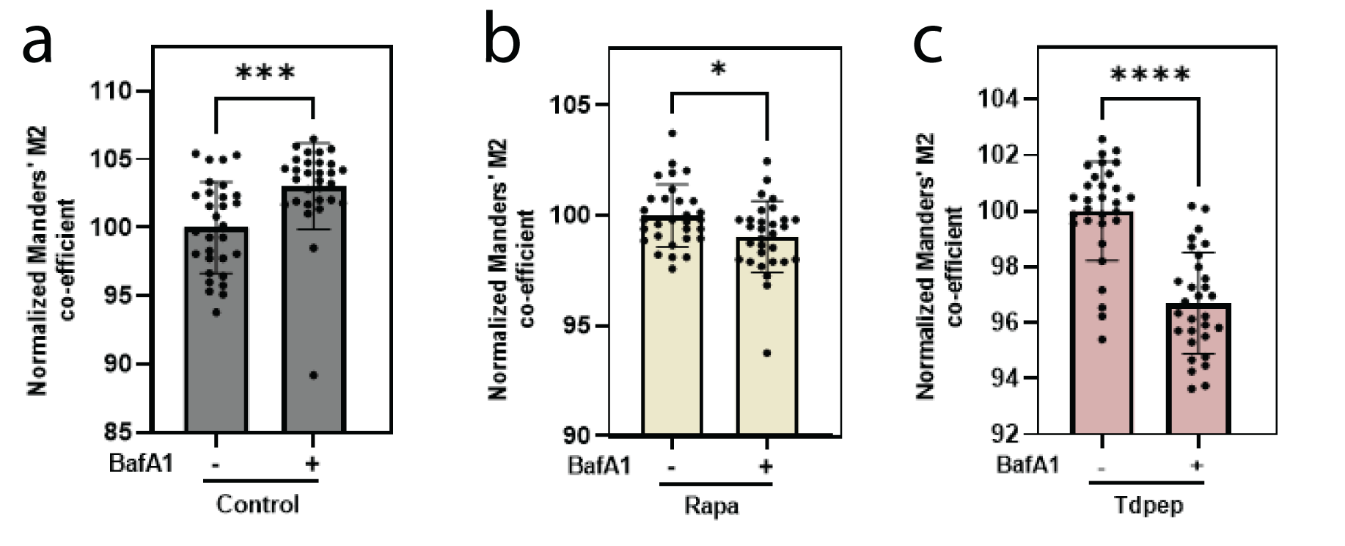

**Figure 2:** a. Manders M2 coefficient of control with Bafilomycin treatment. b. Manders M2 coefficient of rapamycin with Bafilomycin treatment. c. Manders M2 coefficient of Td pep with Bafilomycin treatment

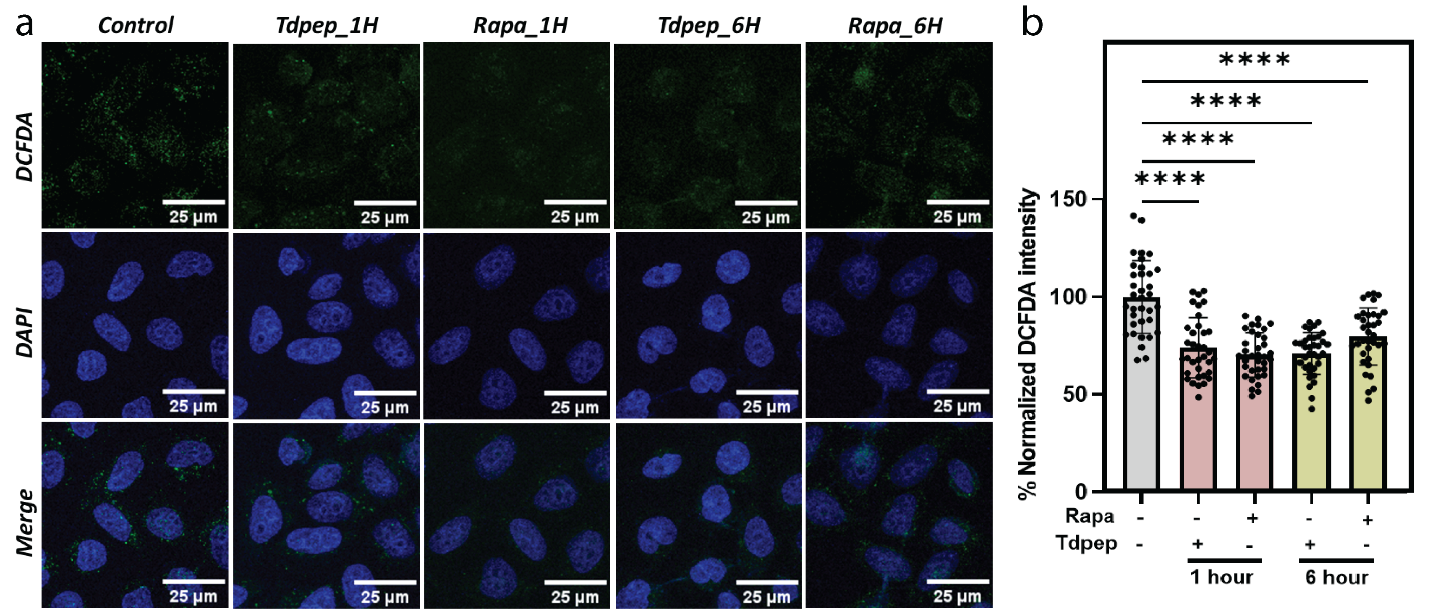

**Figure 3**. a. Confocal images of the ROS analysis with DCF-DA treatment. Scale bar represents 25µm. b. Quantification of the DCF-DA intensities. Histobars represent mean ± SD of 30 cells.

**References:**

1.	Lee EF, Smith NA, Costa TPS da, Meftahi N, Yao S, Harris TJ, et al. Structural insights into BCL2 pro-survival protein interactions with the key autophagy regulator BECN1 following phosphorylation by STK4/MST1. Autophagy. 2019 Jan 9;15(5):785.
